# Repetition Selectively Reconfigures Offset-Defined Temporal Integration in Auditory Cortex

**DOI:** 10.64898/2026.06.01.729053

**Authors:** Xuehui Bao, Peirun Song, Haoxuan Xu, Yuying Zhai, Hangting Ye, Ishrat Mehmood, Nayaab Shahir Pandit, Zhiyi Tu, Lingling Zhang, Xuan Zhao, Xiongjie Yu

## Abstract

Auditory perception requires integrating sound over time, but whether cortical integration windows are fixed or adaptively shaped by recent sensory history remains unclear. Because auditory offset responses are shaped by the temporal structure preceding sound termination, they provide a readout of offset-defined temporal integration. Here, we recorded single-unit activity in awake rat auditory cortex while presenting 512-ms click trains with inter-click intervals of 1, 2, 4, 8 or 16 ms. Each stimulus was repeated ten times, allowing repetition number to index increasing adaptation. Across 271 offset-responsive neurons, repetition strongly compressed the maximal ICI capable of evoking a significant offset response and produced a smaller population-level shift in preferred ICI. These preferred-ICI changes were heterogeneous across neurons, with bidirectional shifts at the single-neuron level. In contrast, the effective temporal window measured with irregular-regular click trains remained stable at the population level, despite neuron-level variability. AAF and A1 further showed distinct repetition-dependent tuning profiles. These findings suggest that adaptation selectively reconfigures ICI-based offset tuning while preserving a population-level local regularity window, revealing flexible and stable components of cortical temporal integration.

## Introduction

In natural listening, meaning is often carried not by isolated acoustic events, but by their temporal arrangement. The rhythm of speech, the flutter of animal vocalizations and the periodic structure of many environmental sounds all require the auditory system to link discrete events into coherent perceptual objects. This operation depends on temporal integration: the accumulation of acoustic information over a finite history before it is transformed into a neural representation^1,2^. Across the brain, temporal receptive windows expand hierarchically, allowing early sensory regions to track rapid fluctuations and higher cortical areas to integrate information over longer timescales^3–6^. In auditory cortex, neurons integrate sound over tens to hundreds of milliseconds, supporting the transformation of sequential acoustic inputs into unified auditory patterns^7,8^. Yet this integration window is unlikely to be fixed. Natural sounds are repetitive, structured and continuously changing, and auditory circuits must constantly adjust to recent sensory history. A central unresolved question is therefore whether adaptation merely changes response strength, or whether it dynamically reshapes the temporal window through which the cortex integrates sound.

Adaptation is a fundamental principle of neural coding. By suppressing redundant responses and reallocating sensitivity toward informative changes, adaptation improves coding efficiency, expands dynamic range and aligns neural representations with current stimulus statistics^9–12^. In the auditory system, repeated stimulation modulates gain, temporal precision and receptive-field structure from subcortical nuclei to cortex^13–17^. Stimulus-specific adaptation is one well-known example, in which responses to common sounds are reduced while responses to rare deviants are preserved^16,18–20^. However, adaptation has usually been examined through firing-rate suppression, frequency selectivity or deviance detection^21–27^. Much less is known about how adaptation affects temporal integration itself. If repeated stimulation changes the timescale over which cortical neurons accumulate acoustic evidence, then adaptation would not simply alter how strongly the brain responds to sound; it would change how the brain segments, groups and interprets sound over time.

The auditory offset response provides a powerful way to address this problem. Although sound offset responses were traditionally regarded as termination signals or rebounds following sound cessation, recent work indicates that they can reflect the temporal structure of the preceding stimulus^28–30^. In primates and humans, cortical offset responses to click trains are strongly shaped by inter-click interval (ICI), stimulus duration and local regularity, revealing a neuronal integration window through which recent acoustic history influences the response at sound termination^31^. Unlike onset responses, which are tightly linked to instantaneous acoustic energy, offset responses provide a delayed readout of what has been accumulated during the preceding sound. This makes them particularly useful for probing the operational timescale of auditory cortex. However, whether this offset-defined integration window is stable or adaptively reconfigured during repeated stimulation remains unknown.

Here we used offset responses to examine how repetition-dependent adaptation shapes temporal integration in awake rat auditory cortex. We presented 512-ms click trains repeatedly with a short silent interval, allowing the first presentation to sample a relatively fresh state and later presentations to sample progressively adapted states. We quantified three offset-derived measures: maximal ICI, which defines the longest temporal interval capable of evoking a significant offset response; preferred ICI, which identifies the interval that most strongly drives the response; and effective temporal window, which estimates how much recent regularity before sound offset is needed to shape the offset signal. This design allowed us to ask whether adaptation uniformly suppresses offset responses or selectively changes specific dimensions of temporal integration. We found that repetition strongly compressed the maximal ICI, modestly shifted the preferred ICI at the population level despite heterogeneous single-neuron changes, and left the effective temporal window largely stable at the population level. These results reveal that adaptation does not simply dampen temporal-integration signals. Instead, it selectively reconfigures ICI-based offset tuning while preserving the local regularity window as a population-level property, suggesting that the auditory cortex adaptively regulates some temporal dimensions while maintaining others as relatively stable reference frames for interpreting ongoing sound.

## Results

### Repetition provides an assay for adaptive temporal integration at sound offset

Offset responses provide a neural readout of temporal integration in auditory cortex. Rather than reflecting only a passive rebound after sound termination, offset activity is shaped by the duration and temporal structure of the preceding stimulus, suggesting that it reports the accumulation of recent acoustic information within a neuronal integration window^28,30,31^. We therefore used offset responses to ask whether repetition-dependent adaptation reshapes temporal integration in auditory cortex.

We recorded extracellular single-unit activity from the auditory cortex of awake, head-fixed Wistar rats (Fig. 1a). Recording sites were assigned to the anterior auditory field (AAF), primary auditory cortex (A1) or ventral auditory field (VAF) using tonotopic maps. Stimuli were 512-ms trains of 0.2-ms clicks with fixed inter-click intervals (ICIs) of 1, 2, 4, 8 or 16 ms (Fig. 1b). For each ICI condition, the same click train was presented ten times consecutively with a 300-ms silent interval between trains, followed by a rest period before the next condition (Fig. 1c). Thus, stimulus number within each ten-presentation sequence provided an operational measure of adaptation strength: the first presentation sampled the least adapted state, whereas later presentations sampled progressively adapted states.

**Figure 1.**
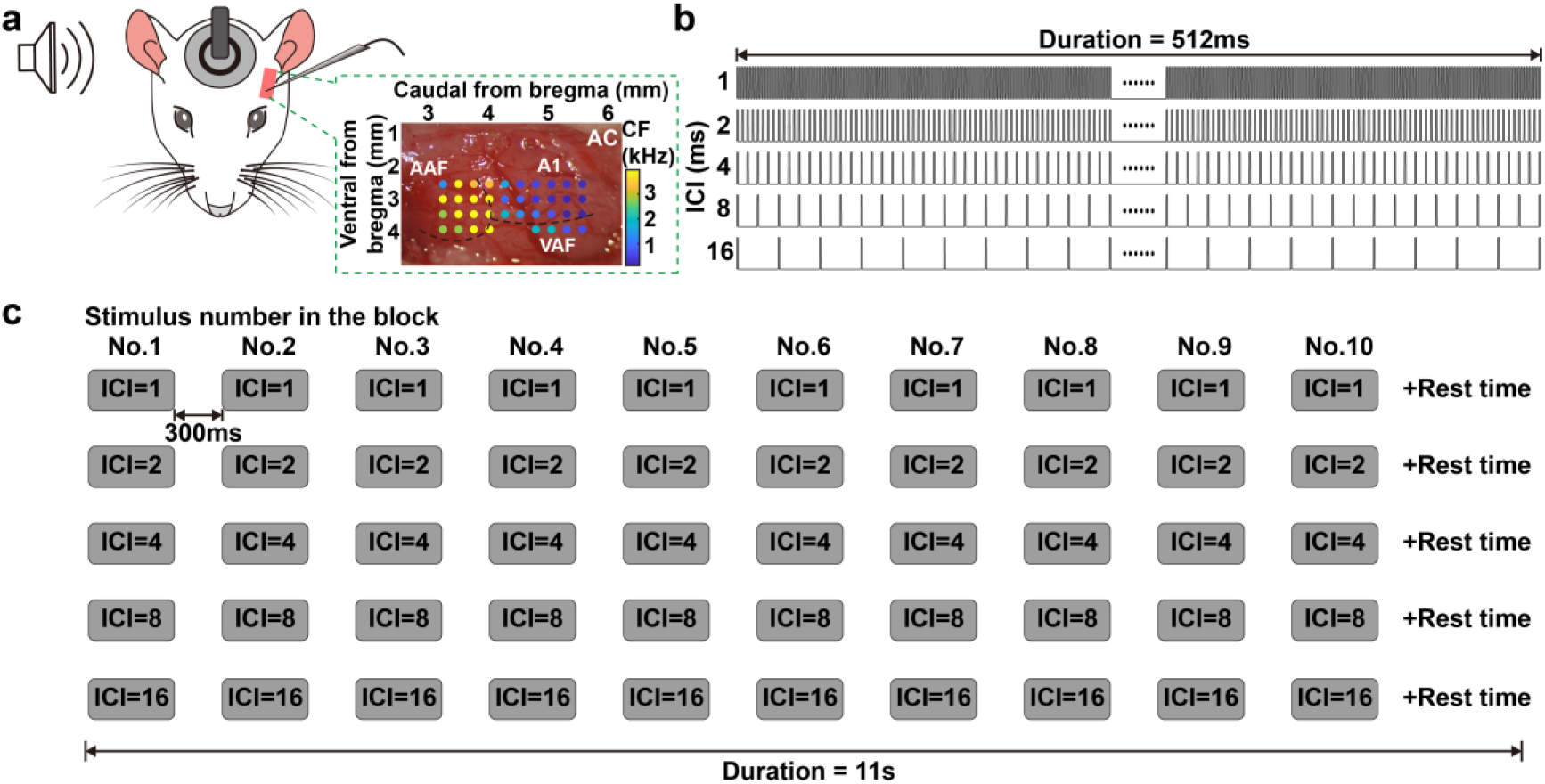
Experimental design for probing repetition-dependent changes in temporal integration in auditory cortex. **(a)** Schematic of extracellular recording in the auditory cortex (AC) of an awake, head-fixed Wistar rat. The inset shows an example tonotopic map used to identify cortical subfields. AAF, anterior auditory field; A1, primary auditory cortex; VAF, ventral auditory field; CF, characteristic frequency. **(b)** Click-train stimuli used in the ICI-tuning experiment. Each stimulus consisted of a 512-ms train of 0.2-ms clicks with a fixed inter-click interval (ICI). ICIs were 1, 2, 4, 8, and 16 ms. **(c)** Repetition paradigm. For each ICI condition, the same 512-ms click train was presented consecutively 10 times with a 300-ms silent interval between consecutive trains, followed by a rest period before the next ICI condition. This repetition structure was used to progressively increase adaptation within each block. The total duration, including the post-block rest period, was 11 s.

Across the auditory cortex, we identified 271 neurons with significant offset responses, defined as firing within 0–80 ms after sound termination that exceeded a late post-offset baseline. We first asked how repetition altered ICI-dependent offset tuning. For each repeated presentation, we quantified two complementary measures: the maximal ICI, defined as the longest click interval that still evoked a significant offset response, and the preferred ICI, defined as the interval that evoked the strongest offset response. These measures allowed us to distinguish changes in the upper temporal boundary of offset responsiveness from changes in the optimal temporal interval driving the offset signal.

### Repetition compresses the maximal ICI of offset responses

We first asked whether repetition altered the temporal range over which click trains could evoke offset responses. In a representative offset-responsive neuron, the first 512-ms click train evoked clear offset-locked firing whose magnitude varied strongly with ICI (Fig. 2a, b). Significant offset responses were present up to an ICI of 8 ms (\**p* < 0.05, *t*-test), whereas the response to 16 ms was not significant (*p* = 0.999, *t*-test); thus, the maximal ICI of this neuron in the first presentation was 8 ms (Fig. 2c). This temporal boundary changed rapidly with repetition. From the second presentation onward, the maximal ICI decreased from 8 ms to 2 ms and remained at this shorter value throughout the sequence (Fig. 2d, e). Because stimulus duration was fixed at 512 ms, this shift reflects a change in ICI-dependent offset tuning rather than a change in stimulus length. This example suggests that repetition can rapidly constrain the temporal range over which preceding click structure supports an offset response.

**Figure 2.**
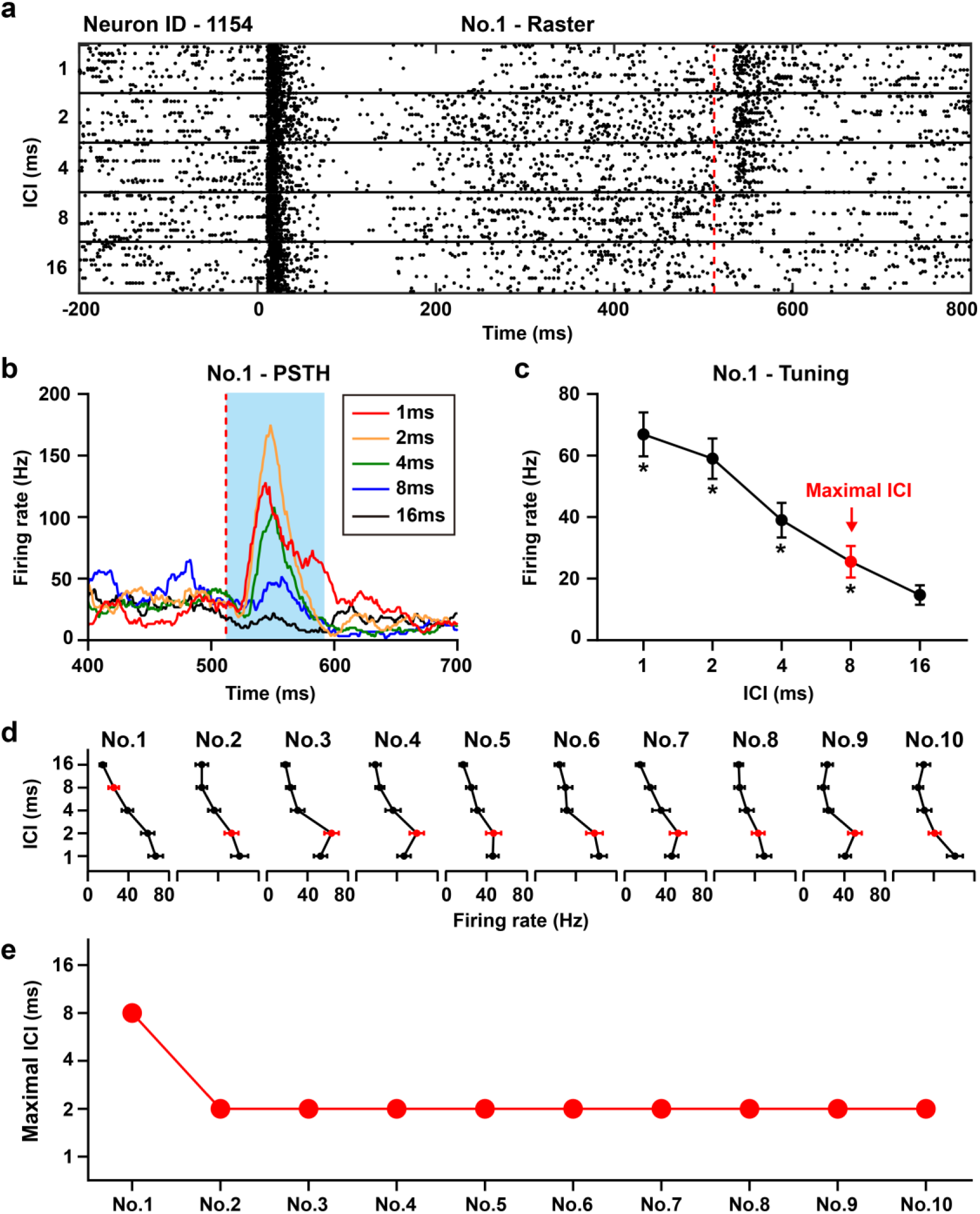
Representative neuron showing repetition-dependent changes in maximal ICI tuning of offset responses. **(a)** Raster plots showing responses of a representative auditory cortical neuron to the first presentation of 512-ms click trains with ICIs of 1, 2, 4, 8 and 16 ms. The red dashed line indicates stimulus offset. **(b)** PSTHs corresponding to the responses shown in **a**, aligned to stimulus offset. The blue shaded region indicates the analysis window used to quantify the offset response. **(c)** Offset firing rate as a function of ICI for block No.1. Error bars indicate standard error (SE). Asterisks indicate significant offset responses. The red marker denotes the maximal ICI, defined as the longest ICI that evoked a significant offset response. **(d)** Offset-response tuning curves across the ten consecutive stimulus blocks. Red markers indicate the maximal ICI for each block. **(e)** Maximal ICI plotted as a function of stimulus block number, showing a rapid reduction from block No.1 to later repeated presentations.

We next tested whether this effect generalized across the population. For each of the 271 offset-responsive neurons, we calculated maximal ICI across the ten repeated presentations. Population distributions shifted with repetition (Fig. 3a). During the first presentation, the mean maximal ICI was 7.61 ms, indicating a sustained but non-monotonic compression of the temporal range capable of driving offset responses. Notably, this adaptation profile was not flat after the initial reduction; instead, maximal ICI fluctuated across repeated presentations, suggesting that repetition-dependent adaptation dynamically modulates the upper temporal boundary of offset responsiveness rather than driving it to a fixed adapted state.

**Figure 3.**
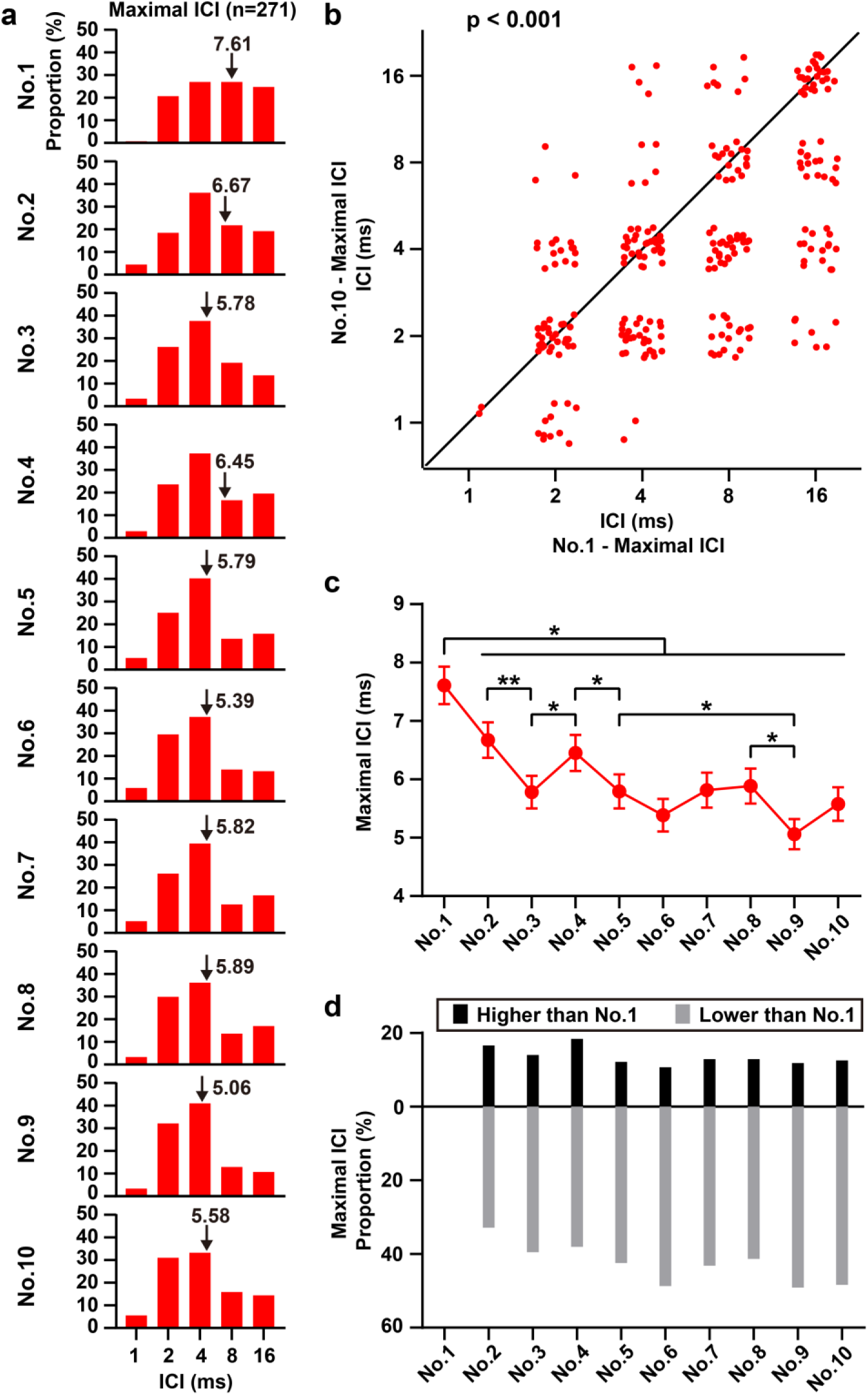
Population-level reduction of maximal ICI across repeated stimulus presentations. **(a)** Distributions of maximal ICI across the 271 offset-responsive neurons for the ten consecutive stimulus presentations. Arrows indicate the population mean for each stimulus number. **(b)** Scatterplot comparing maximal ICI between the first and tenth stimulus presentations. Each point represents one neuron. Points were jittered horizontally and vertically to visualize overlapping observations. Maximal ICI differed significantly between block No.1 and block No.10 (n = 271 neurons, *p* < 0.001, repeated-measures ANOVA with Tukey-Kramer post-hoc test). **(c)** Population dynamics of maximal ICI across the ten repeated presentations. Data are presented as mean ± SE. Asterisks indicate significant differences (\**p* < 0.05, \*\**p* < 0.01, repeated-measures ANOVA with Tukey-Kramer post-hoc test). **(d)** Proportion of neurons showing increased or decreased maximal ICI in each later stimulus presentation relative to block No.1.

A direct comparison between the first and tenth presentations confirmed a significant change in maximal ICI (n = 271, *p* < 0.001, repeated-measures ANOVA with Tukey-Kramer post hoc test; Fig. 3b). Analysis across the full sequence further revealed significant reductions relative to the first presentation for later presentations (Fig. 3c). Individual neurons were heterogeneous: some showed increased maximal ICI, but a larger fraction showed decreased maximal ICI relative to the first presentation (Fig. 3d). Thus, repetition did not impose a uniform shift on every neuron, but biased the population toward shorter maximal ICIs. These results indicate that adaptation selectively constrains the upper temporal boundary of offset responsiveness.

### Preferred ICI is more stable but remains adaptively modulated

Having shown that repetition compressed the maximal ICI, we next asked whether it also changed the ICI that most strongly drove the offset response. Unlike maximal ICI, which measures the upper boundary of significant responsiveness, preferred ICI measures the optimal temporal interval for evoking offset firing.

In the same representative neuron, the strongest offset response during the first presentation occurred at the 1-ms ICI, with progressively weaker responses at longer ICIs (Fig. 4a, b). Thus, although significant offset responses extended up to 8 ms, the preferred ICI was 1 ms. Across the ten repeated presentations, the overall tuning profile remained similar, but the preferred ICI fluctuated between 1 and 2 ms (Fig. 4c, d). This example indicates that preferred ICI is more stable than maximal ICI, but not completely invariant under repetition.

**Figure 4.**
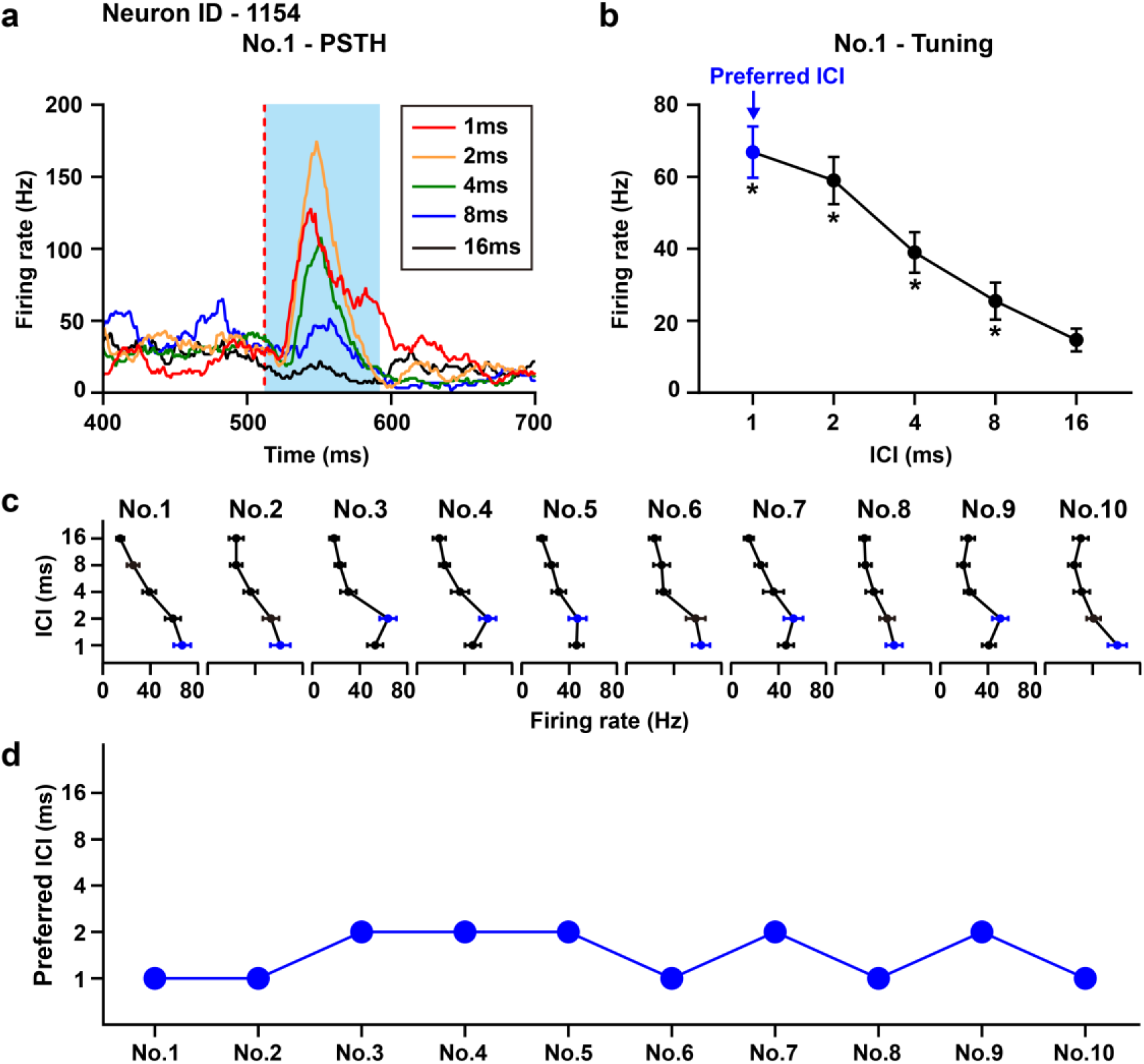
Representative neuron showing preferred ICI tuning of offset responses across repeated stimulus presentations. **(a)** PSTHs showing offset responses of the same representative neuron shown in Figure 2 to the first presentation of 512-ms click trains with ICIs of 1, 2, 4, 8 and 16 ms. The red dashed line indicates stimulus offset. The blue shaded region indicates the analysis window used to quantify the offset response. **(b)** Offset firing rate as a function of ICI for block No.1. Error bars indicate SE. Asterisks indicate significant offset responses. The blue marker denotes the preferred ICI, defined as the ICI that evoked the strongest offset response. **(c)** Offset-response tuning curves across the ten consecutive stimulus blocks. Blue markers indicate the preferred ICI for each block. **(d)** Preferred ICI plotted as a function of stimulus block number, showing modest fluctuations between 1 and 2 ms across repeated presentations.

At the population level, preferred ICIs were concentrated at short intervals, with most neurons preferring 1- or 2-ms click trains across repetitions (Fig. 5a). Nevertheless, the distribution shifted modestly with repetition. The population mean preferred ICI increased from 1.69 ms in the first presentation to higher values in several later presentations, reaching approximately 2.19 ms in the eighth presentation before returning toward shorter values later in the sequence (Fig. 5a, c). Thus, preferred ICI was not driven toward a fixed adapted value; instead, the population trajectory fluctuated across repeated presentations, suggesting dynamic modulation of temporal preference over the course of stimulation (Fig. 5c). A direct comparison between the first and tenth presentations showed a small, marginally significant population-level increase in preferred ICI (n = 271, *p* = 0.047, repeated-measures ANOVA with Tukey-Kramer post hoc test; Fig. 5b). However, individual neurons did not shift uniformly: some neurons increased their preferred ICI, whereas others decreased it, indicating heterogeneous adaptation of the optimal temporal interval at the single-neuron level (Fig. 5b, d). Analysis across the full sequence further revealed significant differences between the first and selected later presentations (Fig. 5c). However, this effect was weaker than the modulation observed for maximal ICI.

**Figure 5.**
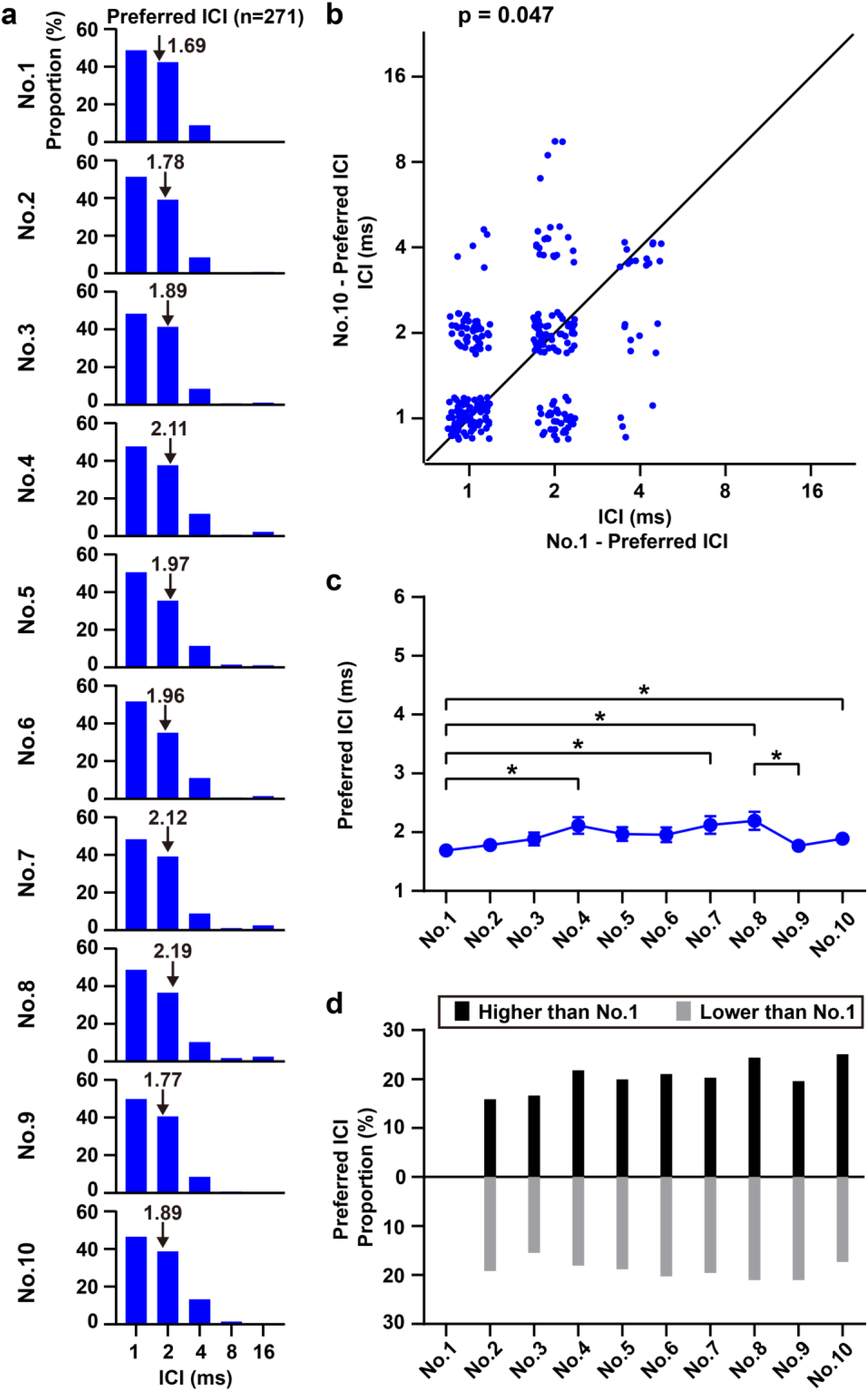
Population dynamics of preferred ICI across repeated stimulus presentations. **(a)** Distributions of preferred ICI across the 271 offset-responsive neurons for the ten consecutive stimulus presentations. Arrows indicate the population mean for each= stimulus number. **(b)** Scatterplot comparing preferred ICI between the first and tenth stimulus presentations. Each point represents one neuron. Points were jittered horizontally and vertically to visualize overlapping observations. Preferred ICI differed significantly between block No.1 and block No.10 (n = 271 neurons, *p* = 0.047, repeated-measures ANOVA with Tukey-Kramer post-hoc test). **(c)** Population dynamics of preferred ICI across the ten repeated presentations. Data are presented as mean ± SE. Asterisks indicate significant differences (\**p* < 0.05, repeated-measures ANOVA with Tukey-Kramer post-hoc test). **(d)** Proportion of neurons showing increased or decreased preferred ICI in each later stimulus presentation relative to block No.1.

Together, the maximal-ICI and preferred-ICI analyses reveal two separable effects of repetition on offset tuning. Repetition strongly compressed the upper temporal boundary capable of evoking a significant offset response, while exerting a weaker and more heterogeneous influence on the optimal interval driving that response. Thus, adaptation primarily constrains the range of offset responsiveness, with a more modest and dynamically fluctuating adjustment of temporal preference.

### Local regularity windows are preserved across repeated stimulation

The ICI-tuning experiments showed that repetition altered two aspects of offset tuning: the upper temporal boundary, captured by maximal ICI, and the optimal temporal interval, captured by preferred ICI. We next asked whether repetition also affected a distinct form of temporal integration: the effective temporal window (ETW) over which recent regularity before sound offset shapes the response. To estimate ETW, we used a separate irregular-regular click-train paradigm with the same total duration of 512 ms and a mean ICI of 4 ms. The control stimulus was fully regular throughout the train. In the irregular-regular stimuli, the initial segment contained randomized ICIs between 1 and 8 ms, whereas the final segment contained regular 4-ms ICIs lasting 128, 64, 32 or 16 ms (Fig. 6a). This design tested how much immediately preceding regularity was required for the offset response to resemble that evoked by a fully regular train.

**Figure 6.**
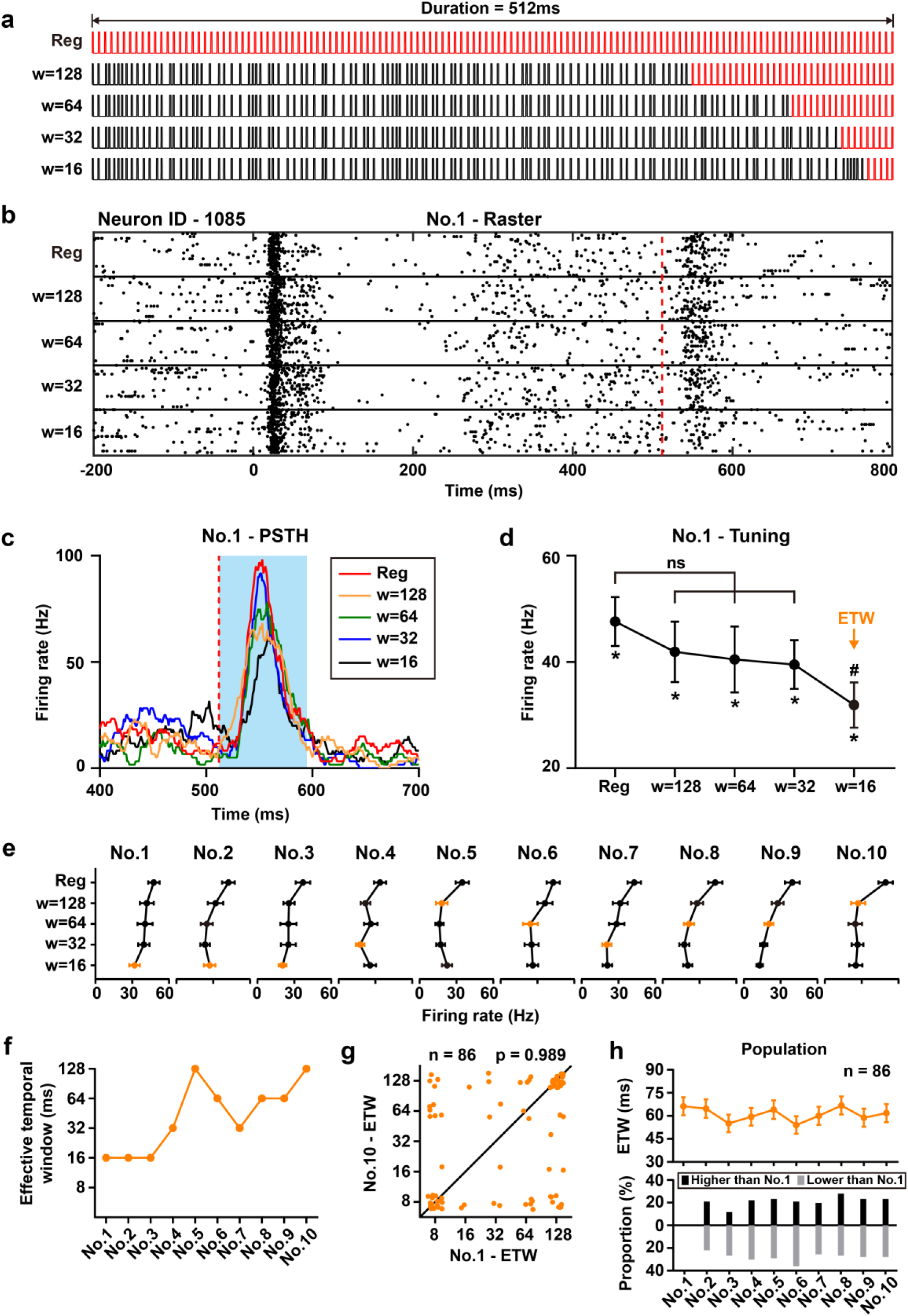
Offset responses to regular and irregular-regular click trains reveal a stable effective temporal window across repeated stimulation. **(a)** Stimulus configuration. Five types of 512-ms click trains were presented. The control stimulus consisted of a fully regular click train with a uniform 4-ms ICI (“Reg”). The other four stimuli consisted of an initial irregular segment with randomized ICIs (mean ICI = 4 ms), followed by a final regular segment with a fixed 4-ms ICI. The duration of the final regular segment was 128, 64, 32, or 16 ms. **(b)** Raster plots showing responses of a representative neuron to the first presentation of the five stimulus types. The red dashed line indicates stimulus offset. **(c)** PSTHs corresponding to the responses shown in **b**, aligned to stimulus offset. The blue shaded region indicates the analysis window used to quantify the offset response. **(d)** Offset firing rate as a function of the final regular-window width. Error bars indicate SE. The 128-, 64-, and 32-ms window conditions did not differ significantly from the regular control. The orange arrow indicates the effective temporal window (ETW), defined as the longest regular-window condition that remained significantly different from the regular control (#*p* < 0.05; ns, not significant). **(e)** Offset-response tuning curves across the ten consecutive stimulus blocks. Orange markers indicate the ETW for each block. **(f)** ETW plotted as a function of stimulus block number for the representative neuron. **(g)** Scatterplot comparing ETW between block No.1 and block No.10. ETW did not differ significantly between the first and tenth stimulus presentations (n = 86 neurons, *p* = 0.989, repeated-measures ANOVA with Tukey-Kramer post-hoc test). **(h)** Population ETW dynamics across the ten repeated stimulus blocks. Top: mean ETW across blocks. Bottom: proportion of neurons showing increased or decreased ETW in each later block relative to block No.1. No significant population-level ETW change was observed across repeated stimulation.

In another representative neuron, the first presentation evoked robust offset responses across all five stimulus types (Fig. 6b, c). Responses to the 128-, 64- and 32-ms regular-window conditions did not differ significantly from the fully regular control, indicating that these durations of recent regularity were sufficient to overcome the influence of the earlier irregular segment. In contrast, the 16-ms regular-window condition remained significantly different from the regular control, indicating that irregular history still influenced the offset response when only the final 16 ms were regular (Fig. 6d). We used this comparison to define the ETW as an operational estimate of the temporal span over which recent stimulus history shaped the offset response.

We then tracked ETW across repeated presentations. In the representative neuron, ETW varied across repetitions, ranging from short windows in early presentations to longer windows in selected later presentations (Fig. 6e, f). This example illustrates that ETW can fluctuate substantially at the single-neuron level. However, these fluctuations did not follow a progressive monotonic trajectory, suggesting that repetition did not drive the local regularity window toward a fixed adapted state in this neuron (Fig. 6f). At the population level, 86 neurons met the inclusion criterion for ETW analysis, requiring significant offset responses to the regular control across all ten presentations. Unlike maximal ICI and preferred ICI, ETW did not differ between the first and tenth presentations (n = 86, *p* = 0.989, repeated measures ANOVA with Tukey-Kramer post hoc test; Fig. 6g). Consistent with the single-neuron variability, individual neurons showed both increases and decreases in ETW between the first and tenth presentations, indicating heterogeneous changes in the regularity window at the neuronal level (Fig. 6g). However, these opposing changes were balanced across neurons, so the population-level ETW remained essentially unchanged. Mean ETW also remained stable across the full sequence, with no significant differences among repetitions (Fig. 6h).

Thus, repetition differentially affected offset-derived temporal metrics. ICI-based tuning, especially maximal ICI, was adaptively modulated, whereas the ETW derived from local regularity remained stable at the population level under the present stimulus regime. This dissociation argues against a uniform degradation of offset responses and instead suggests that adaptation selectively reshapes some dimensions of temporal integration while preserving others at the population scale.

### Offset-defined temporal metrics are organized by cortical field

Finally, we asked whether offset-derived temporal metrics differed across auditory cortical fields. Recording sites were assigned to AAF, A1 and VAF according to tonotopic organization, including orderly characteristic-frequency gradients and reversals in tonotopic progression (Fig. 7a). We focused on AAF and A1, which provided sufficient numbers of offset-responsive neurons for regional comparison.

**Figure 7.**
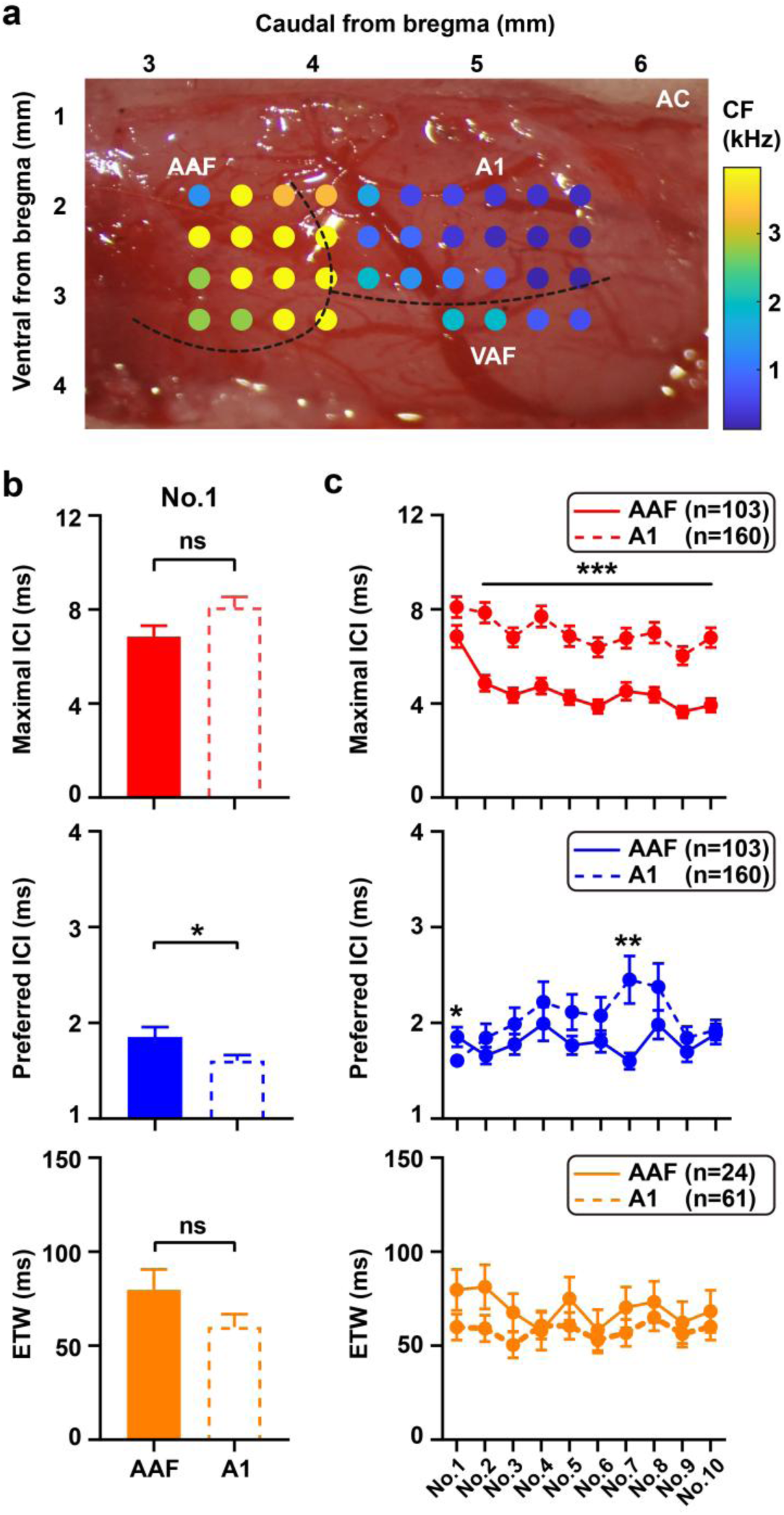
Regional differences in maximal ICI, preferred ICI and effective temporal window between AAF and A1. **(a)** Representative tonotopic map showing the distribution of characteristic frequencies across auditory cortical fields. AAF, anterior auditory field; A1, primary auditory cortex; VAF, ventral auditory field; CF, characteristic frequency. **(b)** Comparison of maximal ICI, preferred ICI and ETW in block No.1 between AAF and A1 neurons. Error bars indicate SE. Statistical significance was assessed using independent samples *t*-tests (\**p* < 0.05; ns, not significant). **(c)** Population dynamics of maximal ICI, preferred ICI and ETW across the ten repeated stimulus blocks in AAF and A1. AAF neurons are shown with solid traces and A1 neurons with dashed traces. Data are presented as mean ± SE. Statistical significance between areas was assessed using independent samples *t*-tests (\**p* < 0.05, \*\**p* < 0.01, \*\*\**p* < 0.001).

During the first presentation, maximal ICI did not differ significantly (*p* = 0.06, independent samples *t*-test) between AAF and A1, indicating that the upper temporal range capable of evoking offset responses was broadly comparable in the least adapted state (Fig. 7b). In contrast, preferred ICI differed significantly (*p* = 0.024, independent samples *t*-test) between the two fields, suggesting regional specialization in the temporal interval that optimally drives offset activity. ETW did not differ significantly (*p* = 0.132, independent samples *t*-test) between AAF and A1, consistent with the population-level stability of this regularity-based measure.

Across the ten repeated presentations, AAF and A1 showed distinct trajectories for ICI-based offset tuning (Fig. 7c). Maximal ICI differed significantly between fields across multiple presentations (\*\*\**p* < 0.001, independent samples *t*-test), indicating that repetition-dependent compression of offset responsiveness was not identical in AAF and A1. Preferred ICI also differed between fields at selected presentations, although these effects were weaker and less consistent than those observed for maximal ICI. By contrast, ETW remained similar between AAF and A1 across repetition, further supporting the relative stability of this measure.

These regional comparisons show that offset-defined temporal integration is not homogeneous across auditory cortex. AAF and A1 both generate offset responses and encode recent temporal structure, but they differ in specific ICI-based tuning properties and in their adaptation-dependent trajectories. Thus, repetition reshapes cortical temporal integration in a structured and regionally organized manner, rather than producing a nonspecific reduction of offset activity across auditory cortex.

## Discussion

Using offset responses as a temporal-integration signal, we found that repeated 512-ms click trains selectively altered offset-defined temporal tuning in awake rat auditory cortex. At the single-neuron level, repetition rapidly reduced the maximal inter-click interval (ICI) capable of evoking a significant offset response (Fig. 2). This effect generalized across the population of 271 offset-responsive neurons, with maximal ICI significantly reduced across repeated presentations (Fig. 3). By contrast, the preferred ICI—the interval that most strongly drove the offset response—was more stable, although still modestly modulated at the population level (Figs. 4 and 5). Importantly, preferred ICI changes were heterogeneous across neurons, with some neurons shifting toward longer ICIs and others toward shorter ICIs. A third measure, the effective temporal window (ETW) derived from regular versus irregular-regular click trains, remained stable at the population level in the subset of neurons eligible for this analysis (Fig. 6). However, individual neurons could show either increases or decreases in ETW, indicating that population-level stability emerged from balanced single-neuron heterogeneity rather than from invariance of every neuron. Finally, regional comparisons showed that AAF and A1 differed in offset-derived temporal metrics and in their repetition-dependent trajectories, particularly for ICI-based measures (Fig. 7).

### Repetition compresses the upper temporal boundary of offset-based integration

The most prominent finding was the rapid reduction of maximal ICI with repetition (Figs. 2 and 3). Because maximal ICI was defined as the longest ICI capable of evoking a significant offset response, this result indicates that repeated stimulation constrained the temporal range over which click trains could still generate an offset signal. In the least adapted state, many neurons responded to click trains with relatively long ICIs; after repetition, significant offset responses were increasingly restricted to shorter ICIs. Thus, repetition appears to compress the upper boundary of offset responsiveness rather than simply reducing firing rate across all conditions.

This finding extends previous work on auditory adaptation. Classical studies have shown that repeated or statistically predictable sounds reduce cortical responses over multiple timescales, reflecting a dynamic adjustment of neural sensitivity to recent acoustic history^9,10,12,14,32^. In many studies, adaptation has been discussed in terms of response gain, stimulus-specific adaptation, or deviance detection^14–16^. Our results suggest that adaptation can also alter a temporal-integration readout: the range of temporal spacing that can still support a sound-offset response. In this sense, the present paradigm shifts the focus from “how much does the neuron respond?” to “over what temporal range can precede acoustic events still be integrated into an offset signal?”

Offset responses are particularly well suited for this question. Although they were historically treated as responses to sound cessation, increasing evidence indicates that they carry information about the preceding sound and are generated by active auditory computations^28–30,33,34^. Our previous work showed that offset responses to click trains in primates and humans depend systematically on ICI and train duration, supporting the concept of a neuronal integration window^31^. The present results build on that framework by showing that this offset-defined temporal range is not fixed, but can be rapidly modified by recent repetition.

A plausible mechanism is that repeated stimulation selectively weakens or gates the synaptic drive supporting offset responses at longer ICIs. Rapid synaptic depression can reshape spectrotemporal tuning in auditory cortex^35^, and adaptive sensory responses can often be modeled as the interaction of multiple filters operating over different timescales^36^. Inhibitory circuits may also contribute. Parvalbumin-positive (PV) and somatostatin-positive (SOM) interneurons exert complementary control over sensory adaptation in auditory cortex^37^, and whole-cell recordings have shown that onset and offset responses can be driven by partly nonoverlapping synaptic inputs^38,39^. In this framework, repeated click trains may preferentially reduce weaker or less synchronized inputs that support offset responses at longer ICIs, leaving only denser click trains capable of producing significant offset firing.

However, the maximal ICI result should be interpreted carefully. Because maximal ICI depends on statistical significance, a reduction in maximal ICI could partly reflect threshold crossing at the long-ICI edge of the tuning curve. The stronger claim is therefore not that every neuron continuously retuned its full ICI-response function, but that repetition shifted the population-level boundary for significant offset responsiveness. This interpretation is supported by two additional observations: preferred ICI changed less strongly than maximal ICI (Figs. 4 and 5), and ETW remained stable under the present assay (Fig. 6). Thus, the data argue against a nonspecific fatigue account and instead support selective modulation of the temporal boundary of offset responsiveness.

### Flexible ICI-based tuning coexists with a stable local regularity window

The comparison between preferred ICI and ETW reveals that offset-defined temporal integration is not a single unitary process. Preferred ICI was concentrated at short intervals and shifted only modestly with repetition at the population level (Figs. 4 and 5). However, this modest population-level shift masked substantial single-neuron heterogeneity, with individual neurons showing either increases or decreases in preferred ICI across repetition. Similarly, ETW remained stable across repeated stimulation in the analyzable subset (Fig. 6), but this stability was a population-level property: individual neurons could show increases or decreases in ETW, and these opposing changes were balanced across the population. This dissociation suggests that different offset-derived metrics capture partly distinct temporal operations and differ in how adaptation is expressed at single-neuron and population scales.

The maximal and preferred ICI measures reflect how the auditory cortex responds to global click spacing across the entire 512-ms train. By contrast, the ETW assay asks how much immediately preceding regularity is sufficient to make an irregular-regular train resemble a fully regular train at sound offset. The population-level stability of ETW therefore suggests that the local regularity window immediately before offset is more resistant to repetition-driven adaptation than the ICI range capable of producing significant offset firing. This does not mean that ETW is fixed in every neuron; rather, it indicates that adaptation-induced ETW changes are not aligned in a single direction across the population under the current stimulus conditions.

Nevertheless, the ETW result should not be overinterpreted as universal invariance. ETW was analyzed only in 86 neurons whose regular-control offset response remained significant across all ten repetitions (Fig. 6), and the terminal regular windows were sampled at four relatively coarse durations. Therefore, the appropriate conclusion is that ETW was stable at the population level under the current stimulus design and inclusion criteria, not that all regularity-based temporal windows are immune to adaptation. Future experiments should sample the regular-window axis more densely, estimate ETW with single-trial or model-based approaches, and test whether stronger adaptation, different mean ICIs, or behavioral engagement can reveal ETW plasticity.

### Regional differences suggest distributed specialization of offset-based temporal integration

The regional comparison in Fig. 7 adds an important organizational dimension. AAF and A1 did not differ significantly in maximal ICI during the first block, but they differed in preferred ICI and showed distinct repetition-dependent trajectories for ICI-based measures. In contrast, ETW remained similar between the two fields. These findings suggest that offset-based temporal integration is not homogeneous across auditory cortex. Instead, different auditory fields may share some temporal operations while differing in how ICI-based tuning is adaptively modulated.

This conclusion is consistent with prior work showing that offset responses are not uniformly distributed across auditory cortical fields. In rodents, AAF has been reported to show stronger or more prevalent offset responses than A1, suggesting field-specific specialization for sound termination processing^29^. Cortical offset responses also contribute to behavioral sound-termination detection^33^, and recent large-scale work indicates that cortical offset activity can provide richer information about preceding sounds than subcortical offset responses^34^. Viewed in this context, the present field differences suggest that regional specialization may concern not only the magnitude or prevalence of offset responses, but also the adaptive dynamics of offset-defined temporal tuning.

Several mechanisms could account for the AAF-A1 differences. One possibility is differential thalamic input: auditory cortical fields receive distinct thalamocortical projections and may inherit different onset/offset balances. A second possibility is differential intracortical amplification. Prior work suggests that AAF may contain stronger intracortical processing of offset responses, whereas A1 may rely more heavily on input-layer responses^29^. A third possibility is that corticothalamic feedback and inhibitory modulation differ between fields. Layer 6 corticothalamic circuits can dynamically adjust sensory gain and tuning precision in auditory cortex and thalamus^40^, and such feedback could plausibly contribute to region-specific adaptation trajectories.

In summary, repeated click trains revealed that offset-defined temporal integration in auditory cortex is neither fixed nor uniformly adapted. Repetition strongly compressed the maximal ICI capable of evoking an offset response, modestly and heterogeneously modulated preferred ICI, and left ETW stable at the population level despite bidirectional changes in individual neurons. These findings suggest that cortical temporal integration is composed of multiple partially dissociable operations: an adaptive boundary for temporal responsiveness, a weakly modulated temporal preference, and a population-stable local regularity window. By using offset responses as a readout of recent acoustic history, this study provides a framework for dissecting how auditory cortex dynamically regulates the temporal structure of sensory processing during repeated sound exposure.

## Methods

### Animals and surgical procedures

Five adult male Wistar rats (250–300 g, 9–12 weeks), free of signs of external ear infection, were selected for this study. The surgical procedures were performed as described previously^41,42^. Sterile surgical techniques were implemented for the implantation of the headpost. Anesthesia was administered using pentobarbital sodium (40 mg/kg, i.p.) along with atropine sulfate (0.05 mg/kg, s.c.) given 15 minutes before the procedure to reduce tracheal secretion. Xylocaine (2%) was applied liberally to the incision site to minimize pain. A head fixation bar was secured to the top of the skull using dental cement and eight titanium screws. Postoperative recovery lasted 7 days, during which anti-edema medications and antibiotics were administered and the weight of animals was monitored daily to ensure their well-being.

After recovery, the left lateral and dorsal surface of the rat skull were exposed. Once cleaned and dried, reference landmarks were established at the bregma and on the left lateral skull for precise electrode positioning. Target area for the electrode recordings was the left auditory cortex. To prevent tissue overgrowth, the skull above the area was thinly coated with dental cement and enclosed with a protective wall. A handheld micro-drill created a small opening (approximately 5 mm × 4 mm) at the designated target site according to a standard rat brain atlas^43^, which was subsequently sealed with brain gel. On the recording day, the dura mater was carefully punctured, and the electrode was vertically inserted into the target area using the established landmarks for guidance.

All procedures in this study received approval from the Animal Subjects Ethics Committees of Zhejiang University (ZJU20210078).

### Extracellular recording

All electrophysiological recordings were conducted in awake, head-fixed rats. In each session, a multi-site linear silicon electrode (32 channels, 2×16 configuration; Institute of Semiconductor, CAS, CN) was utilized, featuring 16 sites on one shank with a 100-μm inter-site spacing. Neural activity was amplified 20000-fold and filtered between 300 Hz and 3000 Hz. Neuronal spike trains were analyzed using offline spike sorting with the Kilosort algorithm, followed by manual curation in Phy.

For the recordings in the AC, a flexible ground wire and a reference wire from the silicon electrode was connected to a titanium screw at the frontal skull area. The probes were carefully inserted vertically through the dura, meticulously avoiding visible blood vessels to minimize tissue damage. The insertion process was facilitated by a single-axis motorized micromanipulator.

White noise stimuli were utilized to confirm the presence of auditory responses. In total, we recorded 271 offset-responsive neurons in the AC.

### Sound stimulation

Experiments were conducted within a sound-proof room. Acoustic stimuli were digitally generated with a computer-controlled Auditory Workstation (RZ6, TDT) at a sampling rate of 100 kHz. These stimuli were delivered through a magnetic speaker (MF1, TDT), the sound intensity for all click trains was uniformly calibrated to 60 dB SPL to ensure consistency across experiments involving different animals. Calibration was performed using a ¼-inch condenser microphone (Brüel & Kjær 4954, Nærum, Denmark) and a PHOTON/RT analyzer (Brüel & Kjær 4954, Nærum, Denmark). All click trains used in the experiments consisted of pulses with a width of 0.2 ms.

To assess the neuronal responses, we conducted two primary measurements: the frequency response area (FRA) and the characteristic frequency (CF). To determine FRA and CF, we presented a series of tones with a duration of 100 ms and a rise-fall time of 5 ms. These tones covered a wide range of frequencies from 0.5 kHz to 47.7 kHz in 26 logarithmic steps. Additionally, we varied the intensity of the tones from 10 to 70 dB SPL in 10 dB steps. Each frequency and intensity combination was repeated five times with 300 ms interstimulus intervals.

In the extracellular recording experiments, we employed two types of click trains to explore the neuronal temporal integration: Firstly, we introduced a series of click trains with a constant duration (512 ms, 10 blocks), systematically varying the ICI ranging from 1 ms to 16 ms. Secondly, to investigate the effective temporal window (ETW), we utilized five types of click trains, each lasting also 512 ms. The control was a standard click train with a consistent ICI of 4 ms (regular). The composition of the other four experimental stimuli was similar: they began with ICIs averaging at 4 ms, varying randomly from 1 ms to 8 ms (irregular), and concluded with a steady ICI of 4 ms (regular), with the number of intervals selected from 32, 16, 8 or 4, which aligned with the window width of the regular part being 128, 64, 32, or 16 ms, respectively.

### Data analysis

All data analysis procedures were done with MATLAB R2024b (MathWorks) and Fieldtrip toolbox^44^. For Spike data, Peristimulus Time Histogram (PSTHs) were generated using a bin width of 20 ms and a step size of 1 ms. The strength of the offset response was quantified as the difference between the firing rate during baseline activity and the firing rate within 0–80 ms window after click train termination. A neuron was classified as exhibiting an offset response when firing activity in this post-stimulus window was significantly greater (*t*-test, *p* < 0.05) than that measured during the late post-offset window (200–300 ms after sound termination).

The ICI that elicited the maximum offset response was defined as the preferred ICI, whereas the maximal ICI was characterized as the upper limit of ICI capable of inducing a significant offset response. Both maximal ICI and preferred ICI were defined only when the offset response was significant. To determine the ETW, we identified the longest window that showed a notable deviation from the control (regular), defining this window as the ETW, which helps quantify the possible size of temporal integration in shaping the offset response. Because neuronal responses exhibited adaptation during repeated stimulation (each stimulus type was presented consecutively 10 times), the offset response in later repetitions could change from significant to non-significant. For ETW analysis, each stimulus type was required to be compared with the control condition, therefore, the offset responses of the control condition had to remain significant across all repetitions (No.1–No.10). Based on the criterion, 86 neurons were selected from the total population of 271 offset-responsive neurons for ETW analysis.

### Auditory cortex parcellation

In this research, the auditory cortex (AC) was functionally subdivided into three major core fields: the primary auditory cortex (A1), anterior auditory field (AAF), and ventral auditory field (VAF), based on their tonotopic organization and neuronal response properties. Multi-site extracellular recordings were performed across the cortical surface, and characteristic frequency (CF) was determined for each recording site using pure-tone stimuli of varying frequencies (0.5–47.7 kHz) and sound pressure levels (10–70 dB SPL). Tonotopic maps were then reconstructed according to the spatial distribution of CFs across penetrations.

A1 and VAF were identified by a continuous posterior-to-anterior tonotopic gradient, in which CFs progressed systematically from low to high frequencies. AAF was defined by a reversal of the tonotopic gradient at the anterior border of A1, exhibiting an opposite frequency progression relative to A1. VAF was located ventral to A1 and showed an independent tonotopic gradient. Cortical borders between subfields were determined according to these frequency reversals and orderly CF transitions.

Only recording sites with clear frequency tuning and stable responses were included for regional classification. Based on these criteria, neurons recorded within the AC were assigned to A1, AAF, or VAF for subsequent analyses, consistent with previous electrophysiological studies in rats^45–47^.

### Statistics

All statistical analyses were performed using MATLAB. Repeated measures ANOVA was used to compare data across multiple groups, followed by Tukey-Kramer post-hoc tests for pairwise comparisons between stimulus blocks. Group-level regional comparisons of preferred ICI, maximal ICI, and ETW between AAF and A1 were assessed using independent samples t-tests. All post-hoc tests were two-tailed, and a p-value of less than 0.05 was considered statistically significant.

## Author contributions

Conceptualization, X.Y.; Methodology, X.Y., X.B., and P.S.; Investigation and Data Curation, X.Y., Y.Z., P.S., H.X., H.Y., X.B., I.M., Z.T., L.Z., and X.Z.; Writing – Original Draft, X.Y., X.B., and P.S.; Writing – Review & Editing, X.Y., Y.Z., and N.S.P.; Funding Acquisition, X.Y. and Y.Z.; Resources, X.Y.; Software, X.B. and P.S.; Validation and Visualization, X.Y., X.B., and P.S.; Supervision, X.Y.

## Acknowledgments

We are grateful to Xiaokai Kou and Sirou Guo for their help with the experiments. This work was supported by STI2030-Major Projects (2022ZD0204600 and 2022ZD0204800) (to X.Y.); National Natural Science Foundation of China 32571216 (to X.Y.), and 32100827 (to Y.Z.).

